# mdciao: Accessible Analysis and Visualization of Molecular Dynamics Simulation Data

**DOI:** 10.1101/2022.07.15.500163

**Authors:** Guillermo Pérez-Hernández, Peter. W. Hildebrand

## Abstract

We present mdciao, an open-source command line tool and Python Application-Programmers-Interface (API) for easy, one-shot analysis and representation of molecular dynamics (MD) simulation data. Building upon the widely used concept of residue-residue contact-frequencies, mdciao offers a wide spectrum of further analysis and representations, enriched with domain specific annotations when possible. It tries offer a user-friendly interface, which simplifies most decisions for non-expert users, while keeping customizability for expert ones. Emphasis has been put into automatically producing annotated, paper-ready figures and tables. Furthermore, seamless on-the-fly query and inclusion of consensus nomenclatures for GPCR, G-proteins, and kinases is made possible through the respective online databases, which allows for bulk selection and comparison across different systems. Finally, the fully documented Python API allows users to include the basic or advanced mdciao functions in their analysis workflows, and provides numerous examples and Jupyter Notebook Tutorials. The source code is published under the GNU Lesser General Public License v3.0 or later and hosted on https://github.com/gph82/mdciao.

## Introduction

Molecular Dynamics (MD) simulations are a widely used tool for the theoretical investigation of the dynamics of (bio)molecular systems with atomic-level detail[1].

In recent years, MD simulation tools have become increasingly user-friendly, and the hardware on which they run has become faster and cheaper[2]. Thus, the challenge that non-expert simulators face shifts slowly from generating the MD trajectory data to analyzing and summarizing it^1^. The depth and scope of this analysis can range from fairly straightforward and intuition-guided to arbitrarily complex and automated[3].

Many software solutions have been produced over the last decades to analyze MD data, offering different degrees of pre-packaged solutions to experts and non-experts. Usually, first and easiest step is to visually inspect the trajectories in 3D using tools such as the popular VMD[4], PyMOL[5], and chimera[6] (among others). While these run locally, other tools like MDsrv[7] or Mol*[8] have recently been put forward to conduct and share 3D MD analysis remotely via web-browser. However, visual inspection is not generally scalable beyond a certain number of trajectories or a certain number of atoms and is often not sufficient to identify or characterize key events.

Hence, very often, the next level of analysis will be offered by these same programs, either via GUI-menus and plugins or programmatically through scripting. Offered are general, community accepted metrics such as root-mean-square-deviation (RMSD), root-mean-square-fluctuation (RMSF), Ramachandran-plots[9], contact-maps, order-parameters, or more specific, user selected geometric values (distances, bond-angles, dihedral angles etc), or interaction types (Hydrogen bonds, salt-bridges, pi-stacking etc). Once arrived at the scripting/programmatic level, tools do not necessarily require a GUI, and can be used, even remotely, directly on the platform where the MD data resides. A very popular example are the analysis tools shipped with the GROMACS MD simulation suite[10], but many other standalone command-line tools provide these (and similar) analysis solutions, e.g. the GetContacts[11] command-line-tool or the popular Python modules MDtraj[12] or MDanalysis[13] All these offer a diverse catalogue of metrics, deliver atomic-level insights, and are available for non-programming experts willing to learn basic scripting.

Finally, data-driven solutions -automated to varying levels-can be considered the next level of analysis, ranging from geometric clustering, to general dimensionality reduction techniques, to more comprehensive, Physics-based modelling like Markov-State-Modelling (e.g. PyEMMA[14], MSMBuilder[15]). When attainable, these models provide a holistic representation of the MD data that is compact, physically accurate, and fully predictive.

However, of particular interest for this paper are tools published as Python modules, in particular those offering an application programming interface (i.e. a Python API) like e.g. MDtraj, MDanalysis, The API enables users to build and combine their analysis workflow with the growing universe of well-documented and well-maintained scientific Python modules[16] (and references therein). Importantly, users can also fully exploit the feature-rich Jupyter Notebook computing environment, which have become a widely popular scientific result-sharing platform[17].

Considering all of the above, mdciao is introduced in this rich software landscape trying to add value with the following ideas:

- Take non-expert users from their MD-data to a set of compact, paper-ready tables and figures in one single shot, while remaining highly customizable for expert users.
- Be able to work with minimal user input.
- Focus on a transparent, transferable, and universal metric that is understandable by experts and non-experts alike: contact-frequencies with hard cutoffs.
- Exploit available consensus nomenclature for bulk selection, manipulation and annotation purposes.
- Place special care on user-friendliness, documentation (inline and online) and tutorials.
- Allow for local computation and representation, i.e. no need to upload data to external platforms.
- Provide expert users a fully-fledged API to integrate mdciao into their workflows without having to leave the Jupyter Notebook platform.

In the following we will outline the principles, inputs, outputs, and features of mdciao in the Design and Implementation section, providing an overview of the command-line-tools in Table 1. Then, for the Results section we use three large composite figures, containing actual inputs and outputs with example data. The examples shown there correspond to one command-line-tool call and to two examples of multiple API calls inside two Jupyter Notebooks. Except for panel d) of Figure 2, no graphics have been edited, i.e. they have been produced and annotated by mdciao automatically. Only some parts of the text outputs have been edited out, denoted as […] and included in the supplementary materials and online documentation.

**Figure 1.**
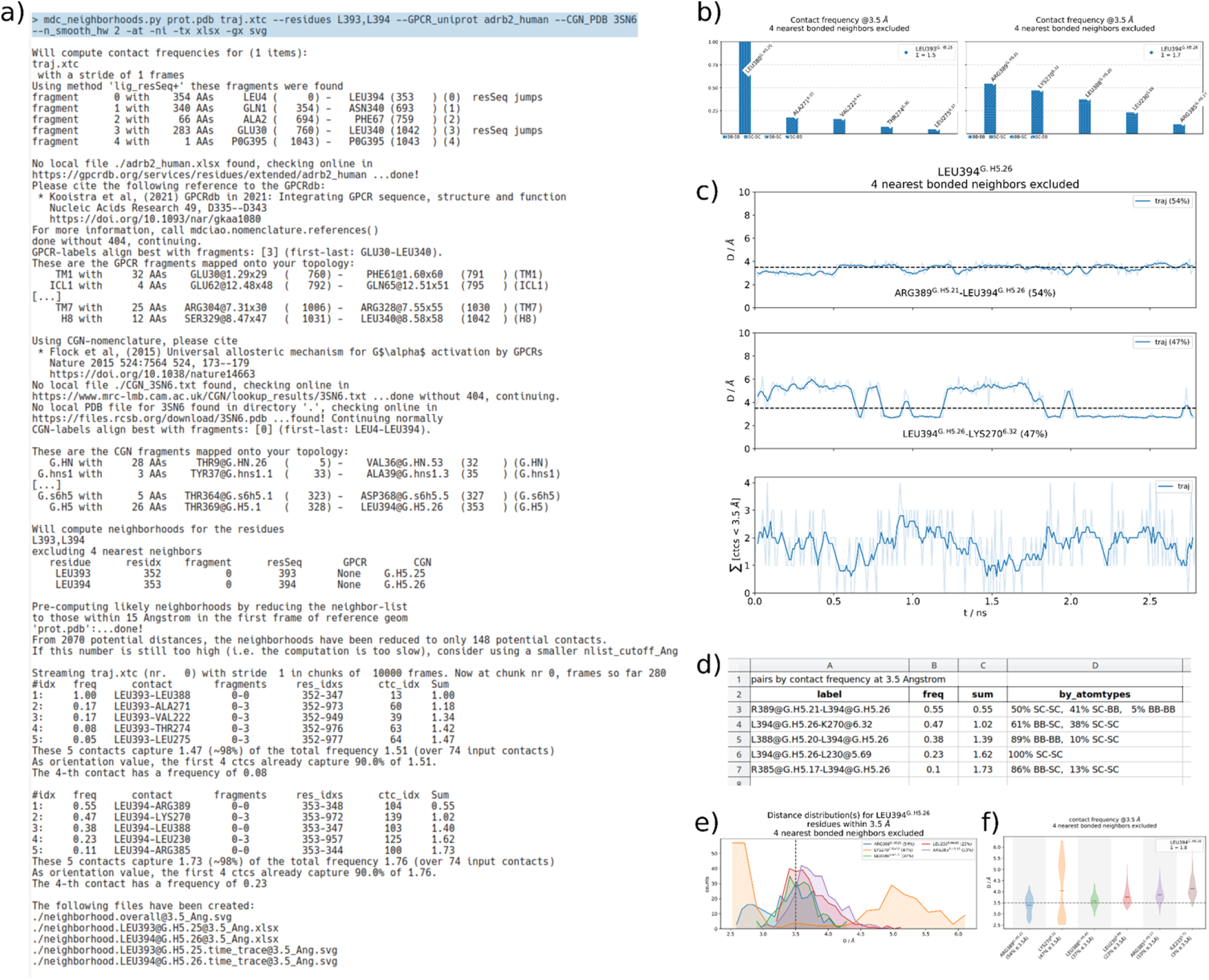
Overview of one `mdc_neighborhoods.py` call from the command line. **a)** Terminal input (on top, shaded) and output, with slight edits denoted as […]. The chosen residues are the C-terminal residues of the Gα-5 helix of the Gαβγ-protein. We can see the fragmentation heuristic recognizing the Gα, Gβ and Gγ subunits (fragment indices 0, 1, and 2, respectively), the β2AR (index 3), and the PG0 ligand (index 4). Then, the uniprot code `adrb2_human` is used to contact the GPCRdb, retrieve consensus labels, align them with the input topology, and map the consensus fragments (TM1 through H8, edited). Analogously, the PDB code 3SN6 is used to retrieve CGN nomenclature. Then, after a computation of likely neighbors, the entire data is scanned for the neighborhoods of LEU393 and LEU394. Once this is done, the individual neighborhoods are reported, and the output files saved and listed. **b)** The file with the contact-frequencies represented as bars, which themselves contain information about interaction types (sidechain or backbone) encoded in their different *hatching* (i.e. the patterns filling the individual bars). Note the consensus labels, which help distinguish between G-protein residues (CGN nomenclature) and receptor residues (GPCR nomenclature with the Ballesteros-Weinstein[25] scheme). **c)** neighborhoodLEU394@ containing the actual time-traces of the residue-residue distances behind the bars in panel a), also annotated with consensus labels and frequency values (three sub-panels have been edited out).The bottom panel of c) also contains the time-trace of the sum over all formed contacts, which oscillates around 1.73 as reported in a). A time-averaging window has been applied to smooth out the fluctuations. **d)** Snapshot of the neighborhood.LEU394@G.H5.2 spreadsheet containing the L394@G.H5.26 neighborhood, numerically specifying the interaction types *hatched* into the frequency bars of b). **f)** Alternative neighborhood representation had the user chosen the option `--report distro`, which plots residue-residue distance-distributions, providing more insight beyond the plain frequency values. **g)** Alternative neighborhood representation had the user chosen the option `--report violins`, which plots residue-residue distance-distributions in violin form, trying to provide a representation as compact as panel b) but as informative as panel f). A full version of these outputs pictures can be found in the supplementary materials, and an online at https://proteinformatics.uni-leipzig.de/mdciao/notebooks/Tutorial.html. Locally, mdciao users can access this CLT example (and others) by invoking the CLT mdc_examples.py (cf. *Table 1*)

**Figure 2.**
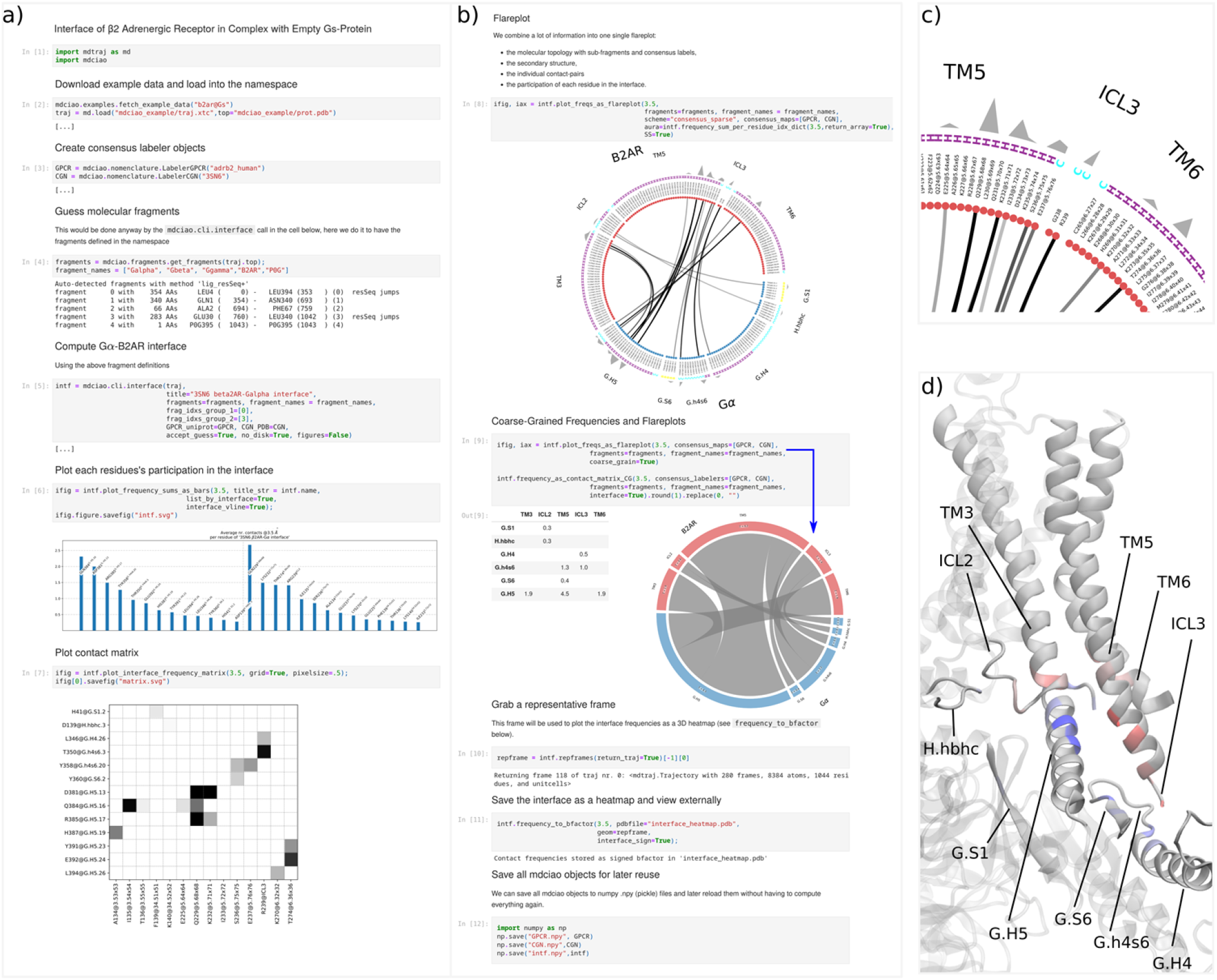
**a)** Example Jupyter notebook illustrating how mdciao can be used in API mode. The notebook consists of 12 cells of Python code (numbered in brackets) and some markdown to provide titles and describe the information flow. Panel a) shows cells [1] to [7], while cells [8] through [12] are shown in b), tiled to the right of a) for a clearer overview. The outputs of cells [3] and [5] have been edited out ([…]) but are analogous to Figure 1A and can be found in the online documentation and the supplementary information. The main computation is the generation of the mdciao object, intf, in cell [5]. It contains the Gα-β2AR-interface contact frequencies and can later be used to generate multiple text, tabular, and graphic reports of the frequencies, distributions, and time-traces. Here, we just highlight a few, namely the per-residue interface-participation (cell [6]), the contact matrix (cell [7]) and the flareplot (cell 8), which, to the best of our knowledge is not implemented in Python as an API anywhere else. As can be seen also in the zoom-in in panel **b)**, the flareplot can integrate many different types of information: the molecular topology with fragments (Gα, β2AR), the consensus subdomains (e.g. TM3 or G.H5), the contact frequencies of the individual residue pairs, the consensus labels of the residues, their secondary structure (letters C for coil, H for Helix and B for β-sheet) and an outer ring (“`aura`”), which can represent any per-residue numerical value. Here, we have plotted each residue’s participation in the interface (i.e., the same bars as in cell [6]), but in principle any other (arbitrary) per-residue quantity could be included into the flareplot, e.g. the sequence conservation degree across a protein class, the root-mean-square-fluctuation (RMSF), the solvent-accessible-surface-area (SASA), the hydrophobicity, or any other informative numerical value that can be imported into the Python namespace. In cell [9], frequencies are coarse-grained to the subdomains, as a table and as chord-diagram. Cell [10] uses `mdciao` to select, within the used MD trajectory data, a frame that is representative of the interface, upon which an interface *heatmap* is added as *bfactor* in cell [11]. Again, this amounts to inserting the bar-heights of cell [6] into the *bfactor* field of the PDB file `interface_heatmap.pdb`, which contains the representative frame. Note that the *bfactor* is signed along interface definitions, meaning Gα-residues get negative *bfactors* and β2AR residues get positive *bfactors*. When reading the PDB-file into any 3D molecular viewer (e.g., in VMD with the blue-gray-red BGrR colormap, shown in panel **d**), the signed *bfactor* highlights the molecular fragments in different colors, Thus, the 2D information of the flareplot can be readily identified, e.g. the middle of the G.H5 (blue) interacting with the tips of TM3, TM5, and ICL3 (red) or the ICL3 interacting lightly with the G.H4 and G.h4s6 subdomains of the Gα (in light blue). This notebook is distributed with mdciao as Manuscript.ipynb and can be run using `mdc_notebooks.py`. A full version of this notebook, with full outputs and high-res pictures can be found in the supplementary materials, and an online version can be found at https://proteinformatics.uni-leipzig.de/mdciao/gallery.html#examples. Locally, mdciao users can access this notebook (and others) by invoking the CLT mdc_notebooks.py (cf. *Table 1*)

**Table 1.**
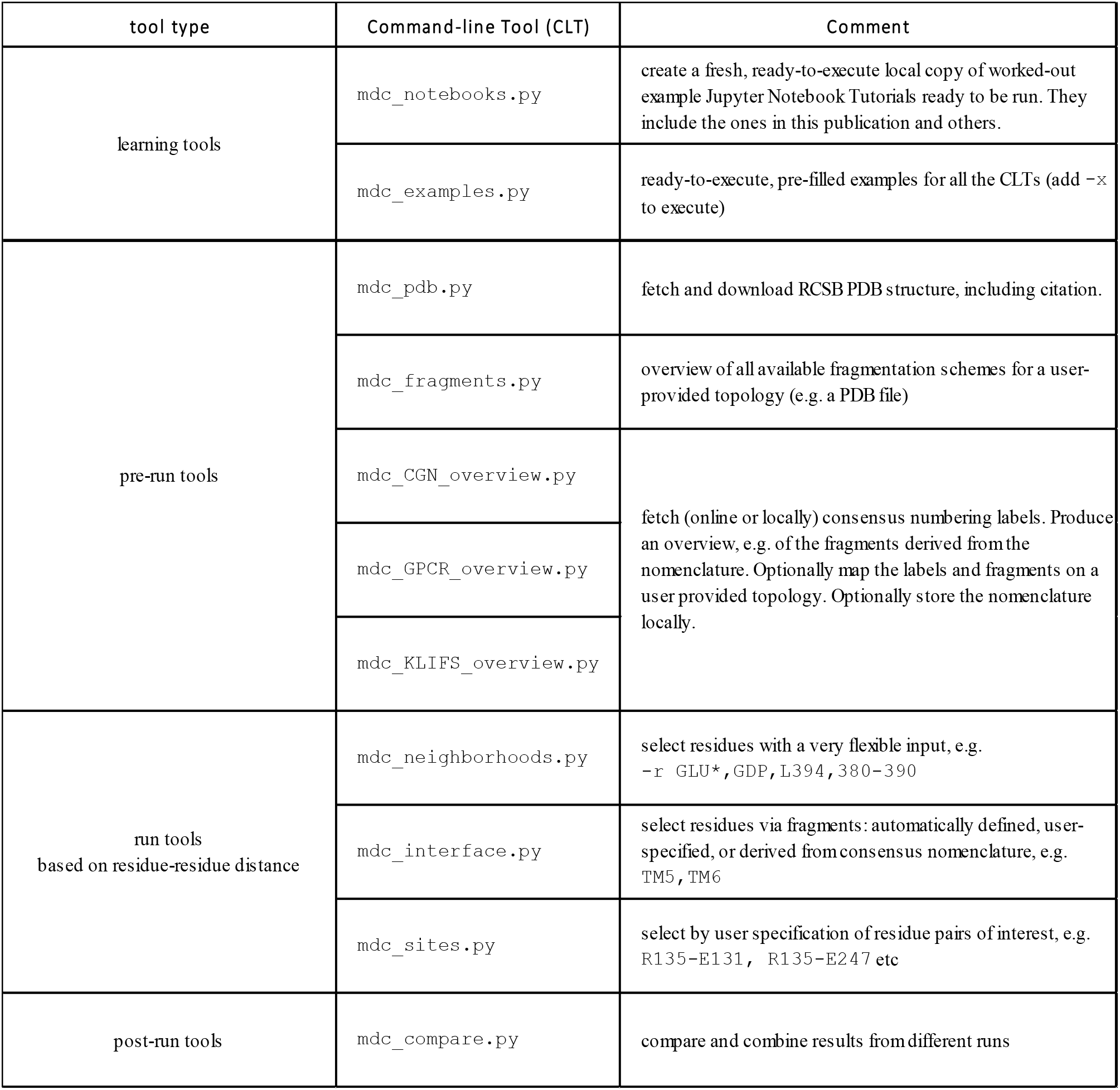
Overview of command-line tools (CLTs) shipped with mdciao. These tools are one-shot tools that take users from basic input to paper-ready figures and tables.

## Design and Implementation

### Basic Principle

At the core of mdciao lies the computation of residue-residue distances, implementing a modified version of the `mdtraj.compute_contacts` method of MDtraj, allowing mdciao to track the atom-types involved in the interactions, e.g. sidechain-sidechain, sidechain-backbone etc. From these distances, contact-frequencies are extracted using a hard distance cutoff of typically 3.5-4.5 Å.

### Input

The needed minimal user input consists of:

- the residues or molecular fragments of interest, such as a ligand, a mutated site etc. two interfacing proteins or subdomains, or arbitrary groups of residues.
- the MD trajectory files to be analyzed.

From this point on, with one command, mdciao automates all fragmentation, labelling, disambiguation, plotting and saving to file. Beyond this minimal input, the user may specify, among many other options (always in one call only). See below the CLT and Jupyter Notebooks for examples of what these options might be.

### Command-line Tools

We present an overview of the command-line tools (CLTs) shipped with mdciao in Table 1. We have divided them into *pre-run, run*, and *post-run* CLTs, with two extra *learning* CLTs to help the user familiarize with mdciao. An example of the inputs and outputs of the CLT `mdc_neighborhoods.py` can be found in Figure 1.

### Fragmentation Heuristics

mdciao implements various heuristics to automatically split the molecular topology into different fragments. These heuristics are independent of the `chain` field of the PDB format, which might not always be correct or be even present, as is the case in the popular.gro-file format. These heuristics use factors such as sequence jumps, presence/absence of bonds, residue names (protein vs non-protein, ion, water) to infer the underlying molecular topology. The so-recovered fragments group residues in meaningful ways, greatly simplifying both the user input and the annotated program output. Example of the fragmentation heuristic being used can be seen in Figure 1, Figure 2 (cell[4]) and in Figure 3 (cell[4]).

**Figure 3.**
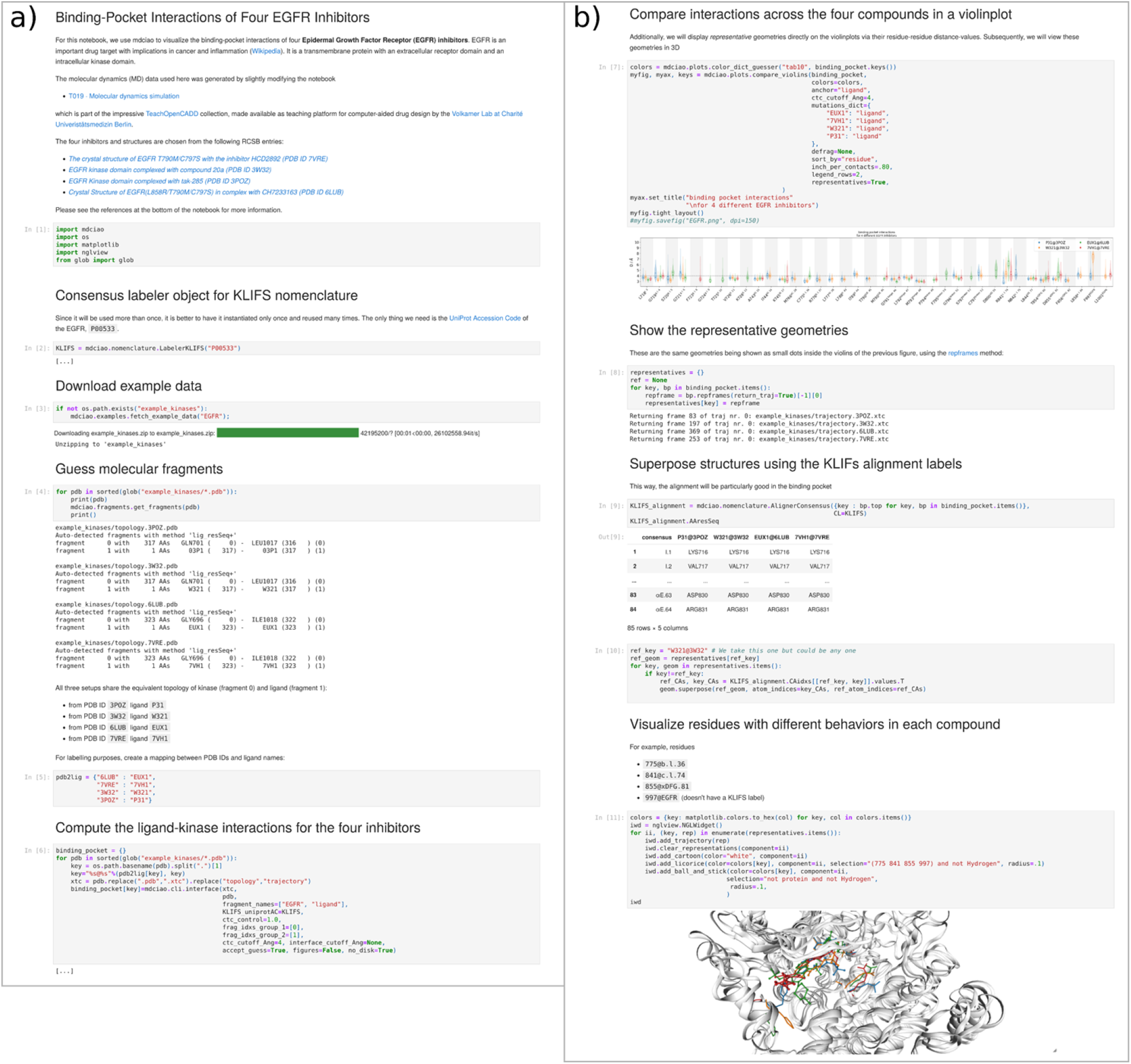
Example Jupyter notebook illustrating how mdciao is used in API mode to compute and compare ligand-kinase interactions for four different inhibitors bound to the Epidermal Growth Factor Receptor (EGFR). The notebook consists of 12 cells of Python code (numbered in brackets) and some markdown to provide titles and describe the information flow. Panel a) shows cells [1] to [6], while cells [6] through [12] are shown in b), tiled to the right of a) for a clearer overview. The outputs of cell [6] has been edited out ([…]) but are analogous to Figure 1A and can be found in the online documentation and the supplementary information. As in Figure 1, the main computation is the generation of the mdciao object `binding_pocket`, using the method `mdciao.cli.interface`, for the four inhibitors of choice EUX1, 7VH1, W321, and P31. As in Figure 2, we generate the nomenclature object on the fly, using the UniProt Accession Code P00533, which is associated to the kinase EGFR. Then, after downloading the example data in [3], we loop over the four datasets and use `mdciao.fragments` to check that the datasets share the same fragmentation, i.e. the kinase and the inhibitor are in the first and second fragments, respectively. Since that is the case, in cell [6] we carry out the same computation four times without any alteration to the API call. Since the goal of the notebook is to compare datasets, we do not show individual plots as in Figure 2, but rather combine all the contact information into one compact violinplot, in cell [7] (please see the SI for a large version of this picture). This plot shows (in vertical) the distributions of the residue-residue distances between the residues of the kinase binding pocket and the three inhibitors (each in one color). The kinase residues are listed along the x-axis and are tagged, with their KLIFS labels. In this plot, one can quickly appreciate individual differences in the binding patterns, in particular when more than one mode (per residue) is present. Furthermore, we use the `representatives` option to superimpose, on top of each violin, a single dot representing the residue-residue distance-value of the representative geometries, which are explained in the next cell. Namely, in [8], we use each `binding_pocket’s repframes`-method to locate (out of the original trajectory data) geometries (=“frames”) representative of the distance distributions shown in the violinplot of [7]. Next, in [9], we exploit the implicit alignment of the KLIFS labels to optimally align, in 3D, the representative frames, which we show in [11]. We do this using the in-notebook 3D molecular viewer nglview. For the 3D plots, we have decided to highlight (in the same color as the violinplots in [7]) the kinase residues C775@b.l.36, P841@c.l.74, D855@xDFG.81, F997@EGFR, which all show different behavior for each inhibitor, as can be seen in the distributions of [7] and in the 3D visualization. A full version of this notebook, with full outputs and high-res pictures can be found in the supplementary materials, and an online version can be found at https://proteinformatics.uni-leipzig.de/mdciao/gallery.html#examples. Locally, mdciao users can access this notebook (and others) by invoking the CLI tool mdc_notebooks.py (cf.*Table 1*)

### Annotations and Consensus Labeling

Whenever possible, all outputs will be annotated using consensus nomenclature labels in texts, tables, and graphics. Currently, the implemented nomenclature databases are the GPCRdb[18] for GPCRs, the Common G-alpha Numbering (CGN[19]) for G-proteins and the KLIFS[20]–[22] for kinases. The user indicates the entry name via UniProt[23] Names, UniProt Accession Codes^2^, or PDB IDs, using either command-line *flags*, e.g. `--GPCR_uniprot adrb2_human` for the CLTs, or as API optional arguments, e.g. `CGN_pdb=‘3SN6’` or `KLIFS_uniprotAC=‘P31751’`. These codes are used to download the consensus nomenclature labels on-the-fly from their respective online databases or, alternatively, to read local files (Excel or plaintext files) which mdciao is able to generate and store for offline use (see Table 1). Subsequently, mdciao maps these labels via pairwise sequence alignment[24] and “tags” residues everywhere in the output with those labels. For example, in the residue-pair R131@3.50-Y391@G.H5.23, an extra layer of information is added succinctly: namely that, in the receptor, R131 is on helix 3, position 50 ([25]) and, on the G-protein, Y391 is on helix 5, position 23 ([19]). Additionally, consensus fragments are automatically inferred and labeled^3^, s.t. mdciao will be aware of exactly what residues (and importantly, what indices) are contained in `TM6` (transmembrane helix 6 for a GPCR) or `G.H5` (helix 5 for a G-protein). These definitions can then, in turn, be used to quickly define interfaces of interest, e.g. for GPCR--G-protein or GPCR—ligand interface. For example, specifying `ICL*` and `G.H*` will compute all contacts between intracellular loops (ICL1, ICL2, ICL3) with the Ras-domain of the G-protein without the user having to define them specifically. This is particularly useful when repeating the same computation for different (but related) systems, where residue indices might have changed and off-by-one errors are likely to happen.

Additionally, mdciao uses the consensus labels can to obtain (multiple) sequence alignments between sequences that share a set of consensus labels but share little sequence identity. Although the four sequences in our EGFR example (Figure 3) are identical, we show how this alignment works in cell [9] and how it is used for optimal 3D superposition in cell [10] and [11].

### Output

While mdciao’s CLTs are running, the *live* terminal output is informing of the different steps taking place. It becomes interactive if user-input is needed, e.g. for disambiguating two equally named residues, and finally produces text reports containing contact frequencies. On top of that, all information is optionally saved to disk as text, spreadsheet, graphic and molecular files. For more details, please see Figure 1.

### API

The Application Programming Interface (API) expands the functionalities of the CLTs and gives the more experienced users programmatic control of mdciao, allowing for the easy inclusion of its methods and classes into arbitrary Python workflows, via `import mdciao`. Crucially, any other, arbitrary Python modules that any user considers of importance for the problem at hand (clustering, time-series analysis, statistical modelling, plotting, formatting) can be used on mdciao’s results without forcing the user to abandon the familiar and powerful (I)Python console or the Jupyter Notebook. This is particularly useful e.g. in Figure 3, where the 3D representation of relevant contacts is carried out “innotebook” using nglviewer[26], and can thus be iteratively fine-tuned while having live access to the data. Finally, all of mdciao’s native objects (classes) can be serialized into NumPy “.npy” files for later use, circumventing the time-intensive computation of a high-number of residue-residue distances.

### Documentation and Built-In Examples

Considerable effort has been invested in making mdciao user-friendly. Firstly, it installs directly with the widely used pip Python manager via `pip install mdciao`. Secondly, `mdc_examples.py` offers new users a catalogue of ready-to-run, pre-packaged CLT-calls that use sample MD data already downloaded at installation. Furthermore, the documentation is extensive and accessible both inline (via the terminal, any integrated development environment (IDE), or the Jupyter Notebook) and online at http://proteinformatics.org/mdciao. There, multiple FAQs, walkthroughs, and Jupyter Notebook Tutorials are presented to showcase most of mdciao’s methods and present potential caveats. Additionally, these notebooks can always be accessed and modified locally in a *sandboxed* way by using the CLT `mdc_notebooks.py` (cf. Table 1).

## Limitations

Using a hard distance cutoff can over- or underrepresent some residue-residue interactions, since not all of these occur at the same residue-residue distance and relative position, e.g. salt-bridges vs. pi-stacking.vs. Hydrogen bonds. Some analysis tools[4], [11], [13] use individual definitions for each interaction type, making the results depend on each of those individual definitions, which may be more or less established or *ad-hoc*.

For sake of simplicity and transferability, mdciao’s results depend parametrically only on one value (the hard cutoff) which is transparently presented in all of the reports. Ultimately, mdciao’s analysis power relies first and foremost on differentiating between frequent and infrequent interactions, and not on slight numerical frequency variations, which will systematically increase or decrease with a given cutoff.

## Results

We present our results in form of overview, multi-panel figures in Figure 1 and Figure 2, and Figure 3, where the extensive captions highlight the information flows and the produced graphics shown in the figure.

Please note that, whereas only a few (of many) use cases have been chosen for this manuscript, readers are highly encouraged to use mdciao’s online tutorials and FAQs to get a full view of the software’s capabilities. It should be noted that mdciao is not a GPCR-specific tool, and can be used with any system, cf. our example notebook on the mutated interface of the SARS-CoV-2 spike protein receptor binding domain (RBD) bound to human angiotensin converting enzyme-related carboypeptidase (ACE2), with data kindly provided by the COVID-19 Molecular Structure and Therapeutics Hub (https://covid.molssi.org/tools/) and the lab of J. Chodera.

## Conclusions

We present a user-friendly command-line tool that produces *one-shot* reports that are paper-ready. It can be incorporated in any Python workflow via its API, and while it analyses MD data locally, it can contact online databases for rich annotation of the results. A variety of plotting functionalities have been implemented to quickly gain insight into the salient features of any MD dataset with little prior knowledge about the system. This tool has already been used in one external publication[27].

## Availability

mdciao is published under the GNU Lesser General Public License v3.0 or later. The source code is hosted on https://github.com/gph82/mdciao, the current stable release is hosted at https://pypi.org/project/mdciao/ and the documentation, including guides and examples can be found at https://proteinformatics.uni-leipzig.de/mdciao. The release used for this manuscript is v.0.0.5.

## Supplementary Information

The entire inputs and outputs of the mdciao calls presented in Figure 1, Figure 2, Figure 3 can be found in the SI.

## Author Contributions

GPH and PWH developed the concept and revised the outputs. GPH designed, wrote, and published the mdciao Python module. All authors wrote the manuscript.

## Acknowledgments

GPH thanks Hossein Batebi and Ramon Guixà-González for their valuable comments and role as beta testers. PWH and GPH thank Sofi Tiwari for her initial contributions to some precursor utilities of mdciao. Additionally, Hossein Batebi, John Chodera, and Andrea Volkamer, are acknowledged for making example data available. This work was supported by Deutsche Forschungsgemeinschaft (German Research Foundation) through SFB1423, project 421152132, subproject C01 and Z04, the Stiftung Charité and the Einstein Center Digital for Future.

## Footnotes

1 Notably, many challenges still remain, like, force-field parametrization of small molecules and overall validity of some physical assumptions of the model, but these are software independent.

2 Please note the difference between UniProt Accession Codes and UniProt Names, as explained here https://www.uniprot.org/help/difference%5Faccession%5Fentryname

3 Please note that this is also valid for kinase consensus fragments, like *linker, hinge*, αD etc.

## Notes

### Competing Interest Statement

The authors have declared no competing interest.

## References

[1] S. A. Hollingsworth and R. O. Dror, “Molecular Dynamics Simulation for All,” Neuron, vol. 99, no. 6. pp. 1129–1143, 2018, doi: 10.1016/j.neuron.2018.08.011.

[2] M. Aldeghi and P. C. Biggin, “Advances in Molecular Simulation,” in Comprehensive Medicinal Chemistry III, vol. 3–8, Elsevier, 2017, pp. 14–33.

[3] A. Glielmo, B. E. Husic, A. Rodriguez, C. Clementi, F. Noé, and A. Laio, “Unsupervised Learning Methods for Molecular Simulation Data,” Chem. Rev., vol. 121, no. 16, pp. 9722–9758, May 2021, doi: 10.1021/acs.chemrev.0c01195.

[4] W. Humphrey, A. Dalke, and K. Schulten, “VMD: visual molecular dynamics.,” J. Mol. Graph., vol. 14, no. 1, pp. 33–8, 27–8, Feb. 1996, [Online]. Available: http://www.ncbi.nlm.nih.gov/pubmed/8744570.

[5] Schrödinger, LLC, “The {PyMOL} Molecular Graphics System, Version~1.8,” Nov. 2015.

[6] E. F. Pettersen et al., “UCSF Chimera--a visualization system for exploratory research and analysis.,” J. Comput. Chem., vol. 25, no. 13, pp. 1605–12, Oct. 2004, doi: 10.1002/jcc.20084.

[7] J. K. S. Tiemann, R. Guixà-González, P. W. Hildebrand, and A. S. Rose, “MDsrv: viewing and sharing molecular dynamics simulations on the web,” Nat. Methods 2017 1412, vol. 14, no. 12, pp. 1123–1124, Dec. 2017, doi: 10.1038/nmeth.4497.

[8] D. Sehnal et al., “Mol* Viewer: modern web app for 3D visualization and analysis of large biomolecular structures,” Nucleic Acids Res., vol. 49, no. W1, pp. W431–W437, Jul. 2021, doi: 10.1093/NAR/GKAB314.

[9] G. N. Ramachandran, C. Ramakrishnan, and V. Sasisekharan, “Stereochemistry of polypeptide chain configurations,” J. Mol. Biol., vol. 7, no. 1, pp. 95–99, Jul. 1963, doi: 10.1016/S0022-2836(63)80023-6.

[10] B. Hess, C. Kutzner, D. van der Spoel, and E. Lindahl, “GROMACS 4: Algorithms for Highly Efficient, Load-Balanced, and Scalable Molecular Simulation,” J. Chem. Theory Comput., vol. 4, no. 3, pp. 435–447, 2008, doi: 10.1021/ct700301q.

[11] “GetContacts,” [Online]. Available: https://getcontacts.github.io/.

[12] R. T. McGibbon et al., “MDTraj: A Modern Open Library for the Analysis of Molecular Dynamics Trajectories,” Biophys. J., vol. 109, no. 8, pp. 1528–1532, Oct. 2015, doi: 10.1016/j.bpj.2015.08.015.

[13] N. Michaud-Agrawal, E. J. Denning, T. B. Woolf, and O. Beckstein, “MDAnalysis: A toolkit for the analysis of molecular dynamics simulations,” J. Comput. Chem., vol. 32, no. 10, pp. 2319–2327, Jul. 2011, doi: 10.1002/jcc.21787.

[14] M. K. Scherer et al., “PyEMMA 2: A Software Package for Estimation, Validation, and Analysis of Markov Models,” J. Chem. Theory Comput., vol. 11, no. 11, pp. 5525–5542, Nov. 2015, doi: 10.1021/acs.jctc.5b00743.

[15] K. A. Beauchamp, G. R. Bowman, T. J. Lane, L. Maibaum, I. S. Haque, and V. S. Pande, “MSMBuilder2: Modeling Conformational Dynamics at the Picosecond to Millisecond Scale.,” J. Chem. Theo. Comput., vol. 7, no. 10, pp. 3412–3419, 2011, doi: 10.1021/ct200463m.

[16] P. Virtanen et al., “SciPy 1.0: fundamental algorithms for scientific computing in Python,” Nat. Methods, vol. 17, no. 3, pp. 261–272, Mar. 2020, doi: 10.1038/s41592-019-0686-2.

[17] J. M. Perkel, “Why Jupyter is data scientists’ computational notebook of choice,” Nature, vol. 563, no. 7729, pp. 145–146, Nov. 2018, doi: 10.1038/d41586-018-07196-1.

[18] A. J. Kooistra et al., “GPCRdb in 2021: integrating GPCR sequence, structure and function,” Nucleic Acids Res., vol. 49, no. D1, pp. D335–D343, Jan. 2021, doi: 10.1093/nar/gkaa1080.

[19] T. Flock et al., “Universal allosteric mechanism for Gα activation by GPCRs,” Nature, vol. 524, no. 7564, pp. 173–179, Aug. 2015, doi: 10.1038/nature14663.

[20] O. P. J. Van Linden, A. J. Kooistra, R. Leurs, I. J. P. De Esch, and C. De Graaf, “KLIFS: A knowledge-based structural database to navigate kinase-ligand interaction space,” J. Med. Chem., vol. 57, no. 2, pp. 249–277, Jan. 2014, doi: 10.1021/JM400378W.

[21] A. J. Kooistra, G. K. Kanev, O. P. J. Van Linden, R. Leurs, I. J. P. De Esch, and C. De Graaf, “KLIFS: a structural kinase-ligand interaction database,” Nucleic Acids Res., vol. 44, no. D1, pp. D365–D371, Jan. 2016, doi: 10.1093/NAR/GKV1082.

[22] G. K. Kanev, C. de Graaf, B. A. Westerman, I. J. P. de Esch, and A. J. Kooistra, “KLIFS: an overhaul after the first 5 years of supporting kinase research,” Nucleic Acids Res., vol. 49, no. D1, pp. D562–D569, Jan. 2021, doi: 10.1093/NAR/GKAA895.

[23] A. Bateman et al., “UniProt: the universal protein knowledgebase in 2021,” Nucleic Acids Res., vol. 49, no.D1, pp. D480–D489, Jan. 2021, doi: 10.1093/nar/gkaa1100.

[24] P. J. A. Cock et al., “Biopython: freely available Python tools for computational molecular biology and bioinformatics,” Bioinformatics, vol. 25, no. 11, pp. 1422–1423, Jun. 2009, doi: 10.1093/bioinformatics/btp163.

[25] J. A. Ballesteros and H. Weinstein, “[19] Integrated methods for the construction of three-dimensional models and computational probing of structure-function relations in G protein-coupled receptors,” 1995, pp. 366–428.

[26] H. Nguyen, D. A. Case, and A. S. Rose, “NGLview–interactive molecular graphics for Jupyter notebooks,” Bioinformatics, vol. 34, no. 7, pp. 1241–1242, Apr. 2018, doi: 10.1093/bioinformatics/btx789.

[27] P. Isaikina et al., “Structural basis of the activation of the CC chemokine receptor 5 by a chemokine agonist,” Sci. Adv., vol. 7, no. 25, p. eabg8685, Jun. 2021, doi: 10.1126/sciadv.abg8685.

